# Revealing the molecular origins of fibrin’s elastomeric properties by in situ X-ray scattering

**DOI:** 10.1101/797464

**Authors:** Bart E. Vos, Cristina Martinez-Torres, Federica Burla, John W. Weisel, Gijsje H. Koenderink

## Abstract

Fibrin is an elastomeric protein forming highly extensible fiber networks that provide the scaffold of blood clots. Here we reveal the molecular mechanisms that explain the large extensibility of fibrin networks by performing *in situ* small angle X-ray scattering measurements while applying a shear deformation. We simultaneously measure shear-induced alignment of the fibers and changes in their axially ordered molecular packing structure. We show that fibrin networks exhibit distinct structural responses that set in consecutively as the shear strain is increased. They exhibit an entropic response at small strains (<5%), followed by progressive fiber alignment (>25% strain) and finally changes in the fiber packing structure at high strain (>100%). Stretching reduces the fiber packing order and slightly increases the axial periodicity, indicative of molecular unfolding. However, the axial periodicity changes only by 0.7%, much less than the 80% length increase of the fibers, indicating that fiber elongation mainly stems from uncoiling of the natively disordered αC-peptide linkers that laterally bond the molecules. Upon removal of the load, the network structure returns to the original isotropic state, but the fiber structure becomes more ordered and adopts a smaller packing periodicity compared to the original state. We conclude that the hierarchical packing structure of fibrin fibers, with built-in disorder, makes the fibers extensible and allows for mechanical annealing. Our results provide a basis for interpreting the molecular basis of haemostatic and thrombotic disorders associated with clotting and provide inspiration to design resilient bio-mimicking materials.

---

In Nature, many examples are found of elastomeric materials made of protein fibers. Prominent elastomeric proteins in the human body are the intermediate filament cytoskeleton of cells^1^, elastin and fibronectin in tissues^2,3^, and fibrin networks in blood clots and wounds^4^. These proteins all form filaments that can reversibly stretch up to tensile strains of around 150% and resist strains of several hundred percent without breaking^5,6^. Two main molecular mechanisms have been identified to accommodate large deformations. The first is the unfolding of ordered domains, for example in silk fibroin and muscle myosin^7,8^. The second is the uncoiling of flexible, disordered domains, as happens in titin^9^. Among the various elastomeric protein systems found in Nature, fibrin networks are a particularly interesting example, since fibrin fibers extend through a combination of these two mechanisms^5^. Fibrin fibers are built from rod-shaped fibrin monomers that contain folded domains that can stretch by unfolding^10^ as well as natively unstructured domains that can stretch entropically^11^. This stretchability allows fibrin clots to stiffen by at least two orders of magnitude before rupture^12^, which provides blood clots with an enormous mechanical resilience against the shear forces exerted by flowing blood and the traction forces exerted by cells^13,14^.

Given the importance of fibrin elasticity for haemostasis and thrombosis^15^, there has been a long-standing interest in the molecular mechanisms by which fibrin achieves its unique elastomeric behaviour. The complex hierarchical architecture of fibrin is generally thought to be a key factor^16^. The soluble precursor protein fibrinogen is a symmetric trinodular molecule that is 45 nm in length, made up of two sets of three polypeptide chains denoted Aα, Bβ and γ^17,18^. Both sets of chains form a coiled-coil connector from the distal (D)-nodules to the center, where they converge in the E-nodule. Near the ends of the molecule, the Aα chains expose long and flexible carboxy-terminal regions known as the αC-regions^19,20^. Thrombin enzymatically converts fibrinogen to fibrin, initiating self-assembly of monomeric fibrin into protofibrils comprised of two linear strands of fibrin monomers that are precisely half-staggered due to specific interactions between neighbouring D- and E-nodules^21^. Once the protofibrils reach a critical length, they bundle into thick (typically 100 nm diameter) fibers which are strengthened by enzymatic crosslinking by Factor XIIIa (FXIIIa)^22^. Along their axis, fibrin fibers exhibit the same half-staggered packing periodicity as protofibrils^23–25^ but in cross-section, evidence from light and X-ray scattering suggests that protofibrils are packed in a fractal, partially crystalline array^26–28^. The fibers in turn form a random network structure that is branched and crosslinked to varying degrees, depending on the solution conditions, temperature, and protein concentrations^29,30^.

The hierarchical architecture of fibrin networks provides several possible mechanisms that may contribute to fibrin’s elastomeric properties. On the molecular scale, single-molecule experiments and molecular dynamics simulations have shown that fibrin monomers can lengthen substantially through forced unfolding of the C-terminal γ-chain nodule and the coiled-coils^10,31^. Simulations predict that stretching converts the molecular conformation of the connector regions from α-helical to β-sheet^32,33^, providing a possible mechanism to explain the nonlinear stiffening of fibers observed at large tensile strains^34^. At the fiber scale, an additional mechanism for elongation emerges as a consequence of the intrinsically disordered αC-regions, which form long and flexible tethers connecting the more rigid protofibrils^35^. The most direct evidence for this mechanism comes from stretching experiments on fibers assembled from fibrinogen from different animal species, showing that the fiber extensibility correlates with the length of the αC-region^36^. Further evidence comes from observations of the recoil dynamics of stretched fibrin fibers, which was shown to be faster than expected from a mechanism involving protein refolding^37^. Orientationally disordered networks of fibrin fibers have an additional mechanism by which they can accommodate strain, namely by the alignment of fibers towards the direction of principal strain^38–40^.

Experimentally it is very challenging to pinpoint the contribution of each of these mechanisms to the mechanical response of a fibrin network. Recently, a few studies have started to address this challenge by performing *in situ* measurements of structural changes in mechanically deformed fibrin networks. Vibrational spectroscopy has revealed evidence of monomer unfolding (α-helical to β-sheet conversion) in stretched fibrin networks^41,42^, Small Angle X-ray Scattering (SAXS) has provided evidence of monomer unfolding based on changes in the axial packing periodicity of fibers within stretched clots^39^ visible through a shift of the Bragg peak^26^, and confocal imaging has shown that fibers reorient and can irreversibly lengthen upon repeated straining^43^. Altogether, there is convincing evidence that molecular unfolding occurs at macroscopic tensile strains above 100%, but it remains unclear to what extent other mechanisms (in particular fiber reorientation and stretching of the unstructured αC-regions) also contribute to the macroscopic elasticity, and how this depends on strain^44^.

Here we present measurements that provide direct evidence of the structural mechanisms that explain the elastomeric properties and nonlinear elasticity of fibrin through *in situ* SAXS experiments on fibrin networks subject to a macroscopic shear deformation. To this end, we combine a shear rheometer equipped with a concentric cylinder (Couette) measurement cell that is transparent to X-rays with a SAXS setup, using a synchrotron source to enable dynamic measurements (Fig. 1a, b). We show that the SAXS scattering patterns provide simultaneous access to shear-induced changes in alignment of the fibrin fibers, through the anisotropy in the scattering patterns, and to changes in the molecular packing structure of the fibers, through the position and width of the Bragg peak corresponding to the periodic axial packing order. We find that increasing levels of shear strain induce distinct changes in network stiffness that coincide with a sequence of structural responses on different scales. Our measurements provide direct support for recent models of fibrin elasticity^45^ that propose a combination of entropic (network-scale) elasticity at small strains and enthalpic (fiber-scale) elasticity at high strains. Our measurements furthermore strongly support earlier proposals^35,46^ that reversible stretching of the αC regions plays an important role in fiber elongation.

**Figure 1.**
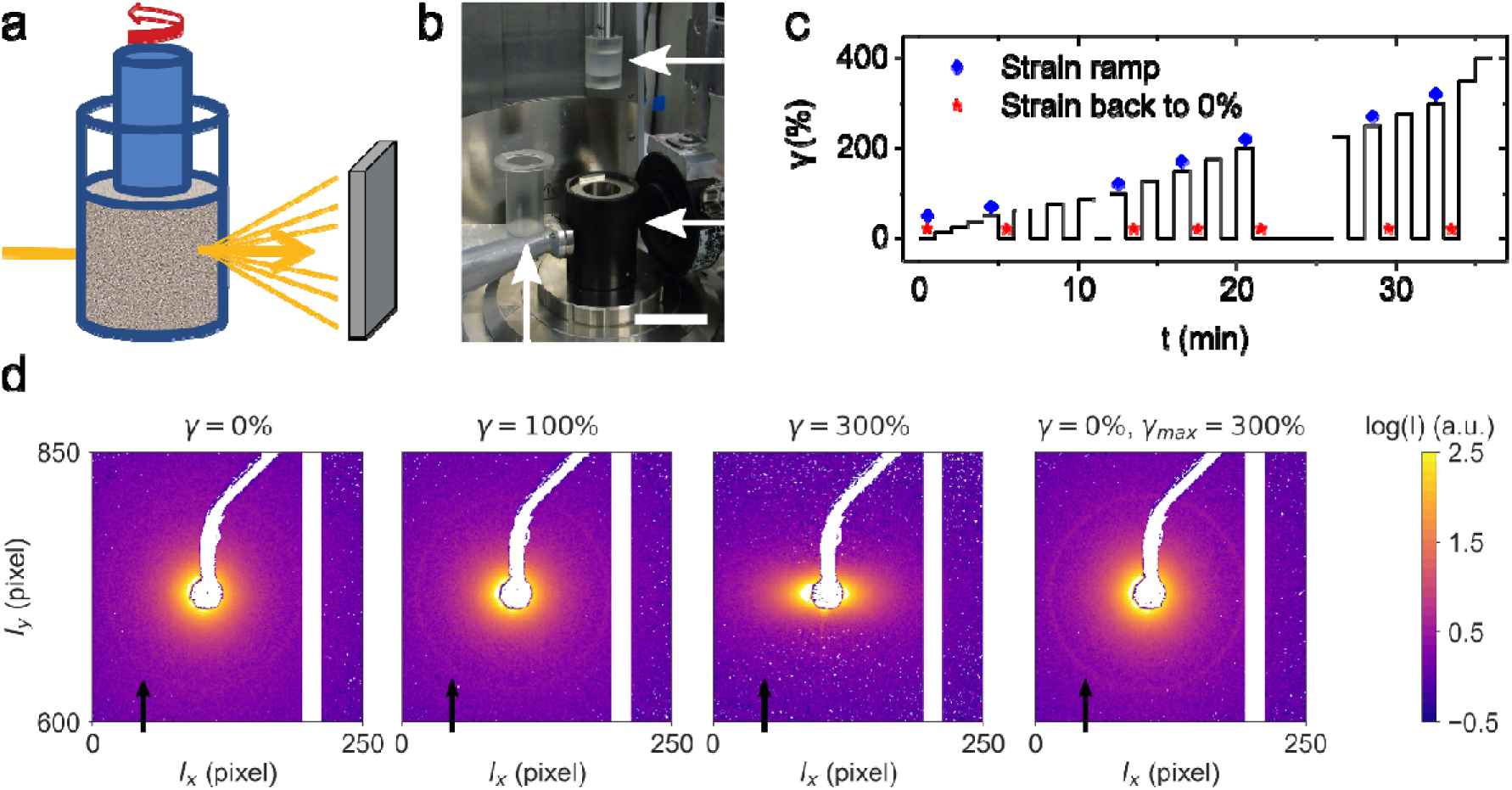
*in situ* SAXS measurements coupled with shear rheology on fibrin networks. (a) Schematic representation of the experimental setup, consisting of a rheometer with a transparent Couette cell placed in the X-ray beam in front of a detector. The sample is sheared by rotating the inner cylinder of the Couette cell while the outer cylinder remains stationary. Unscattered light is stopped by the beamstop while scattered light reaches the detector. (b) Photograph of the setup, showing the inner cylinder attached to the rheometer (top right arrow) and the outer cylinder placed next to its holder (bottom right arrow), which is connected to a Peltier element for temperature control. The scale bar is approximately 5 cm. (c) The strain protocol for shear experiments with stepwise increases in shear strain, interspersed with intervals where the strain is brought back to zero. The blue diamonds refer to the strain levels shown in Fig. 2c, while the red stars correspond to Fig. 4a. (d) Background-subtracted scattering images measured for three different levels of applied shear strain (see legends) and upon return to zero strain on the same 4 mg/ml fibrin gel. The first order scattering ring is (dimly) visible around the beamstop, as indicated by the black arrow. Data are shown on a logarithmic scale (colour bar on the right).

## Results

### Shear-induced structural changes at the network and fiber level

In order to probe the effects of shear strain on fibrin network structure, we combined rheology experiments on fibrin networks in a Couette cell with *in situ* SAXS measurements (Fig. 1a-b). We subjected the networks to a stepwise increase in shear strain γ interspersed with intervals where the strain was brought back to zero to test for reversibility (Fig. 1c). Fig. 1d shows the background-subtracted scattering images of a 4 mg/ml fibrin gel at strains of 0%, 100% and 300%, and the same network returned to zero strain. We observe a high-intensity ring around the beamstop, corresponding to the Bragg peak that originates from the half-staggered packing order of fibrin fibers^47^. Under quiescent conditions, the ring is circular, indicating that the network is initially isotropic. As strain increases, the scattering pattern becomes anisotropic, elongating along the horizontal direction and indicating that fibers align along the shear direction. In addition, increasing the strain leads to a progressive decrease of the intensity of the Bragg peak, indicating a decrease in the molecular packing order of the fibers. Upon return to zero strain, the scattering pattern regains a circular shape indicative of a return to an isotropic arrangement. Moreover, the intensity of the ring increases beyond its original level, indicating that the fibers regain an ordered packing structure with enhanced order compared to the original state.

To investigate the effect of network deformation on the molecular packing structure of the fibrin fibers more closely, we performed a quantitative analysis of the strain-dependent changes in the Bragg scattering peak (Fig. 2b). To this end, we radially integrated the scattering patterns, taking the beam stop as the center (Fig. 2b). The spectra show an overall decrease in scattering intensity as a function of wavevector *q*. At small *q* (i.e. large length scales), the intensity decreases as a power-law in *q*, because scattering occurs off the fractal network structure^48,49^. At high *q*, scattering instead occurs from the internal structures of the fibers. In this regime, we observe a clear peak around *q* = 0.3 nm^−1^, corresponding to the first order Bragg diffraction expected from the half-staggered packing periodicity of 22.5 nm. The shape and position of the peak provide information on the packing structure of the fibers. At zero strain, we also recognize weak higher-order diffraction peaks corresponding to the third and fourth order reflections (indicated by the black arrows). The second order reflection (expected at q = 0.56 nm^−1^, indicated by the grey arrow) is suppressed, consistent with prior SAXS measurements^23^.

**Figure 2.**
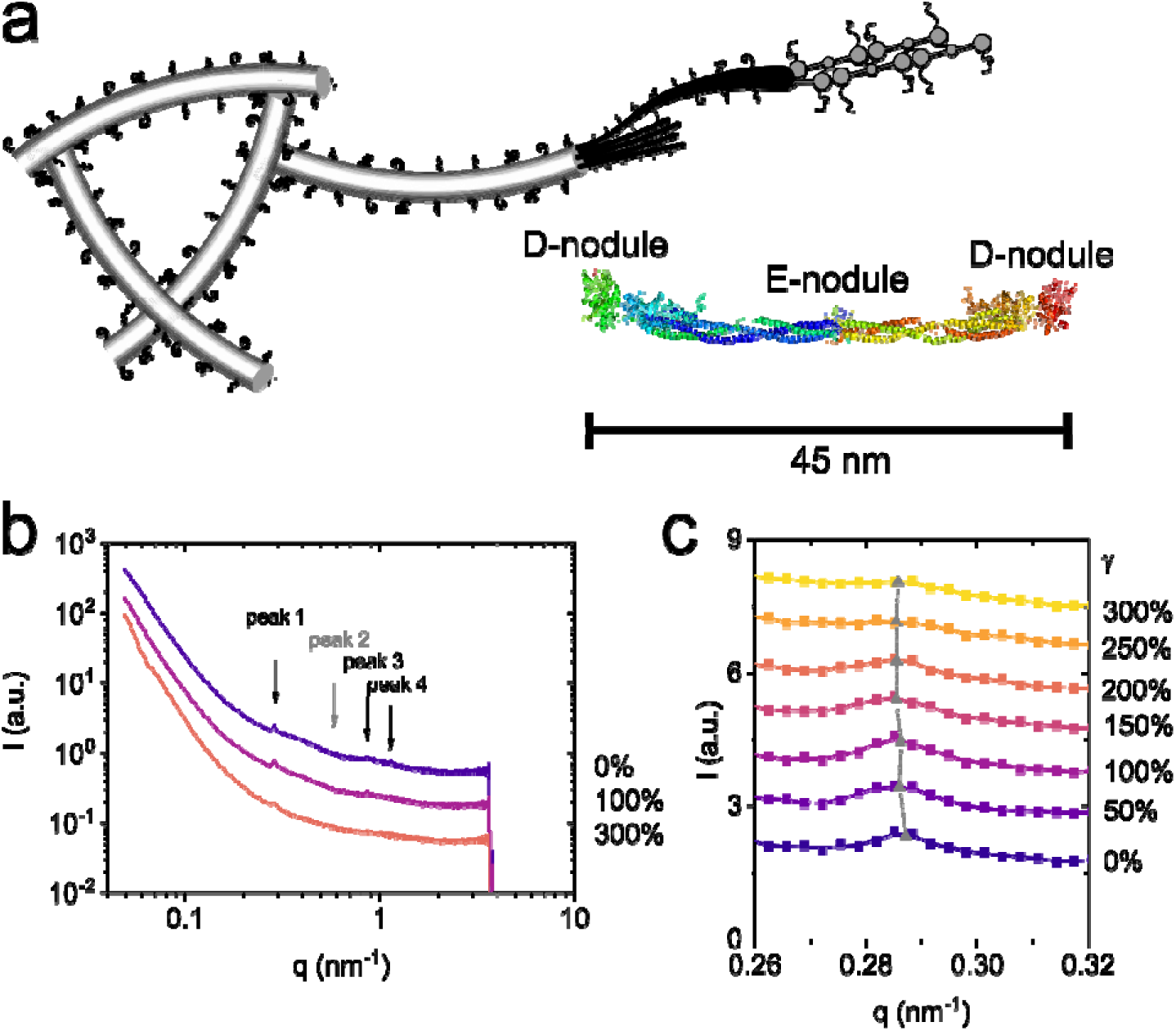
Shear-induced changes in the fiber packing structure. (a) Schematic representation of the hierarchical structure of fibrin, progressively zooming in from the network level, to the fiber level, to the protofibril level, and ultimately the monomer level. The molecular structure of the monomer is shown in the bottom right corner, with the distal D-nodules, central E-nodule, and α-helical coiled-coil connector regions. The crystal structure is taken from Ref. ^17^. Note that the intrinsically disordered αC-regions do not show up in the crystal structure. (b) The radially integrated intensities of the scattering patterns (see Fig. 1d) at different shear strains, showing intensity as a function of scattering vector *q*. The arrows indicate the position of the Bragg scattering peaks. (c) Zoomed-in regions of the scattering spectra (squares) with the overall intensity convoluted with a Gaussian to enable peak-fitting of the first order Bragg peak (lines). The grey squares indicate the peak position *q*_peak 1_, obtained from fitting the radially integrated scattering curve. For clarity, curves obtained at increasing applied shear strain (legend on the right) have been shifted along the y-axis.

To infer the strain-dependence of the packing structure, we applied a fitting procedure to extract the first order Bragg diffraction peak. After subtraction of the overall decrease of intensity with q, we fitted the peaks to a Gaussian function (Supplementary Figure S5). It can be clearly seen that with increasing shear strain, the peaks become less sharp and the position shifts to slightly lower *q*-values (Fig. 2c). To quantify these changes, we used three parameters: the peak position *q*_*peak* 1_, which we associate with the axial repeat distance; the full-width half-maximum (FWHM) of the peak, which reflects the distribution of sizes in the repeat distance and perhaps also instrumental noise^50^; and the peak height *h*, which we associate with the degree of order of the periodic packing of the monomers.

Fig. 3a shows how the position of the Bragg peak develops with strain, obtained from integration in the quadrants of the scattering spectra that are oriented in the direction of strain (‘q1’ in Supplementary Figure S1). Up to shear strains of nearly 100% we observed no significant change in the peak position, but beyond 100% strain the peak position progressively shifted with increasing strain, from *q*_*peak 1*_ = 0.287 nm^−1^ at rest towards *q*_*peak 1*_ = 0.285 nm^−1^ at the highest strain levels. This observation indicates a slight molecular elongation. This peak shift continued until the network suddenly ruptured at a strain around 250%. Rupture was evident from the mechanical response by a sudden drop in stiffness to a vanishingly small value caused by the disappearance of a percolating network (Fig. 3d), and also from the SAXS patterns by a return of the shape from anisotropic to isotropic. Fig. 3b shows the evolution of the corresponding peak height for the same samples. The peak height is constant up to strains of ~50% and shows a slight upturn before the strain reaches 100% and before the peak position starts to noticeably shift, and progressively decreases with strain beyond this peak. This observation indicates that there is a progressive disruption of the spatial order.

**Figure 3.**
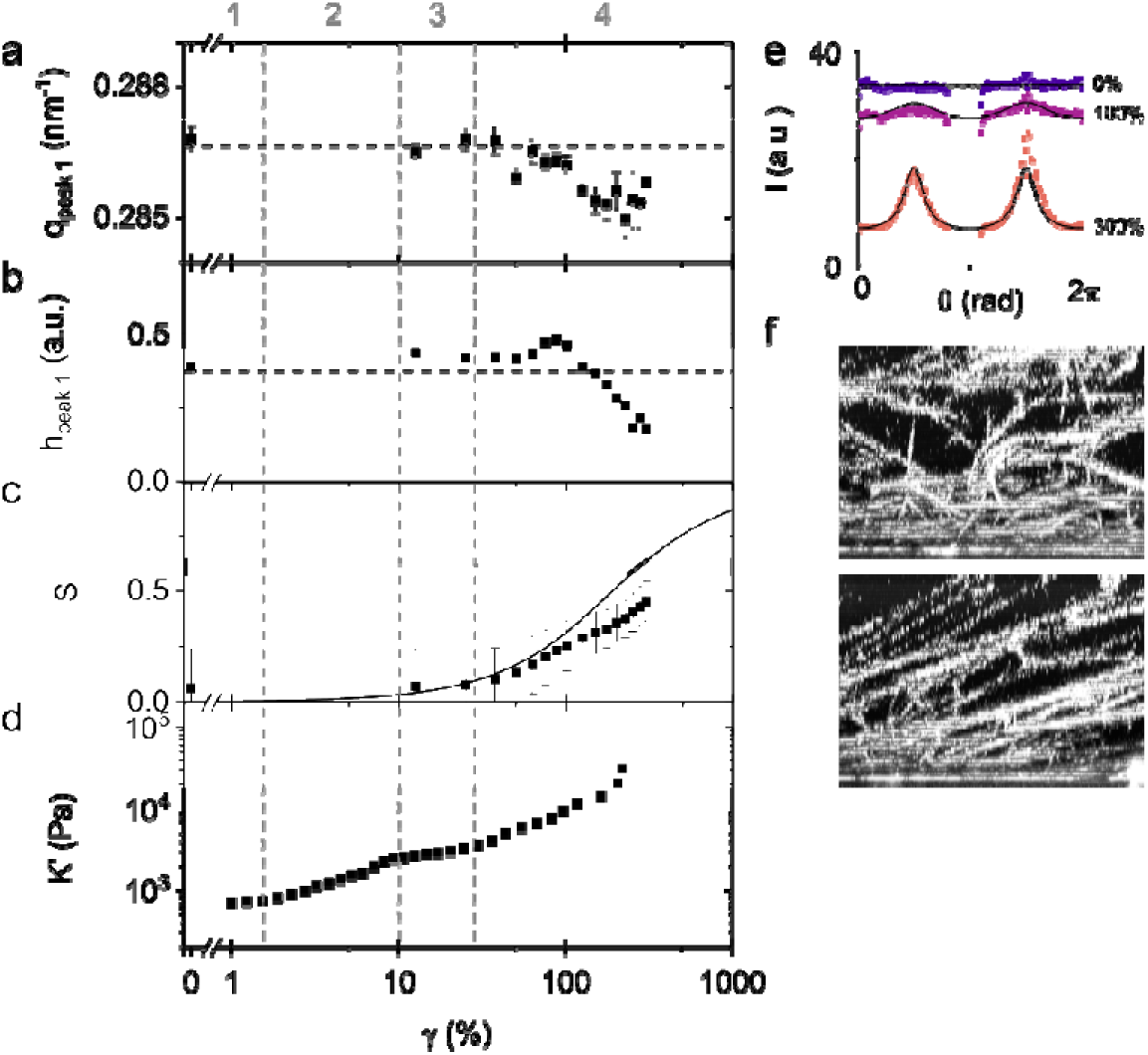
Correlation between strain-dependent elasticity and structural changes across scales. The position (a) and height (b) of the first scattering peak as a function of γ. The dashed horizontal lines indicate the value of the peak position and height of the original, unstrained fibrin network. Error bars are the standard deviation between two independent measurements. (c) The nematic order parameter *S*, calculated from the azimuthally integrated scattering intensity, as a function of γ for two independently prepared samples. The solid line shows the model prediction for the case of an affinely deforming, initially random ensemble of rigid rods. (d) The differential elastic modulus *K*’ of a 4 mg/ml fibrin gel as a function of γ. We observe 4 distinct mechanical regimes that are indicated by numbers (top) and separated by dashed vertical lines (see main text). (e) Azimuthally integrated intensities, showing intensity as a function of angle θ. The spectrum is flat at zero strain, as expected for an isotropic sample, and develops peaks indicative of fiber alignment with applied strain. Fiber alignment is fully reversible upon removal of the strain. The black curves show the predicted scattering pattern for an initially random ensemble of rigid rods, undergoing an affine shear deformation. Around θ = π data is missing due to the beamstop. For clarity, the curves have been shifted along the *y*-axis. (f) Confocal images at zero strain (top) and 200% strain (bottom) confirm fiber alignment.

The maximal shift of the Bragg peak position reached at 250% applied shear strain, just before network breakage, corresponds to a length increase of only 0.3 nm, or 0.7% of the fibrin monomer length. This discrepancy between the large network strain on the one hand and the tiny monomer strain on the other hand raises the question of what the tensile strain felt at the fiber level is. The strain at the fiber level will depend on how uniformly the macroscopic strain is distributed across the network. For a continuum elastic solid under shear, the local strain is everywhere identical and equal to the macroscopic strain. However, filamentous networks, including fibrin, are known to be prone to nonaffine behaviour^51–53^, although the degree of nonaffinity is difficult to predict *a priori*. Any nonaffinity would be expected to delay the onset of alignment and lower the alignment compared to the affine prediction. To test whether the deformation is affine, we therefore compared our measurements with the alignment predicted in the limit of an affine deformation. We analysed the evolution of the fiber orientations with increasing shear strain from the anisotropy of the scattering patterns and compared it to the affine prediction. We azimuthally integrated the scattered intensity over a *q*-range corresponding to length scales between 49.7 and 102 nm. Here the lower limit is set such that the Bragg peak is excluded, and the upper limit is set by the lowest *q*-value we can access. As shown in Fig. 3e, the intensity is angle-independent at 0% strain and develops peaks at angles of ¼π and ¾π upon the application of a shear strain. These peaks are well fit by an orientational distribution function for stiff fibers^54^ (solid lines).

From these fits, we computed the nematic order parameter *S*, following the approach in Ref.^55^. This number can range between 0 for fully isotropic fibrous networks, and 1 for fully aligned fibrous networks. As shown in Fig. 3c, the nematic order parameter starts out at values close to zero, consistent with an initially random network structure, and smoothly increases to a value of around 0.5 at a shear strain of 250%. For comparison, we plot the expected evolution of the order parameter with increasing shear strain in case of an affinely deforming random distribution of non-interacting fibers (solid line). We find that the measured nematic order parameter closely follows the affine prediction up to applied shear strains of 100%. At higher strains, it increases less rapidly than the affine prediction, likely because the fibers are constrained by the presence of junctions, visible in confocal images taken under shear (Fig. 3f).

Given that the shear-induced fiber alignment to a good approximation follows the affine limit, we can estimate the elongation of the fibers from the applied shear strain using the affine model. We thus estimate that at an applied shear strain of 250%, the tensile strain on fibers oriented in the direction of strain is 83%. This strain is still well below the maximum strain of ~250% that fibrin fibers can sustain without breaking^6^. Strikingly, the total fiber elongation of 83% is two orders of magnitude larger than the 0.7% increase in monomer length inferred from the shift of the Bragg peak. This observation makes a fiber elongation mechanism mediated by simultaneous monomer unfolding unlikely (see Discussion).

### Correlating SAXS and rheology measurements

Concurrently with the SAXS measurement, we also measured the dependence of the elastic modulus of the network on the applied shear strain (Fig. 3d). With increasing strain, we observe four distinct mechanical regimes (demarcated by vertical dashed lines): (1) a linear elastic regime (γ < 2%), (2) a first strain-stiffening regime (2% < γ < 10%), (3) a short plateau regime where the modulus is again strain-independent (10% < γ < 25%) and (4) a second strain-stiffening regime that continues until the network breaks (25% < γ < 250%). This striking multi-step stiffening response has also been reported in previous rheological studies of fibrin networks, where it was interpreted in the context of theoretical models for semiflexible polymer networks^12,56^. It was proposed that the elastic modulus at small strain is entropic in origin because fibrin fibers are semiflexible and therefore store some excess length in thermal bending undulations^12,56^. Small shear strains will straighten out the filaments and pull out the entropic slack. Since the filaments are rather stiff, strain-stiffening sets in already at strains of a few percent. At larger strains, the elastic modulus is expected to be enthalpic in origin, because the filament backbones experience tensile strain. Based on the SAXS measurements, we are now able to test this interpretation directly by correlating the rheological data with the observed changes in network and fiber structure.

In the linear elastic regime and the first strain-stiffening regime, we found no measurable changes in the position and height of the Bragg peak (Fig. 3a-b), nor in fiber alignment (Fig. 3c). This observation is consistent with the notion that the elasticity is entropic in origin at these small levels of strain, in the sense that no structural changes are expected on the molecular scale as long as the fiber backbone does not experience any tensile strain. Once the strain reached 25%, we simultaneously observed the onset of renewed network strain-stiffening (Fig. 3d) and the onset of fiber alignment, indicated by an increase in the nematic order parameter (Fig. 3c). This observation is consistent with the notion of enthalpic elasticity, whereby progressive filament recruitment along the direction of principal strain causes network stiffening^38–40^. Although the filament backbones experience tensile strain, the position of the Bragg peak nevertheless remained unchanged until the strain reached ~100% strain, suggesting there is not yet any molecular elongation. We did observe a slight increase in the height of the Bragg peak at strains just below 100%, probably because fiber alignment gives rise to a sharper Bragg peak. Once the strain went beyond 100%, the Bragg peak position shifted to shorter *q* while the peak height decreased, indicating that the monomers experience sufficient tensile strain such that they elongate. The shift of the Bragg peak coincided with an apparent transition of the strain-stiffening response to a somewhat steeper increase of the stiffness with strain right before network rupture.

### Reversibility of shear-induced structural changes

To gain more insight in the mechanisms that accommodate strain in sheared fibrin networks, we tested the reversibility in the structural changes described above upon return to the initial unstrained configuration. At the network level, we probed the anisotropy in the scattering pattern after return to zero strain. Since the fit procedure to obtain the nematic order parameter is too sensitive to noise at low anisotropy, we employed an alternative measure of anisotropy at low strain. Specifically, we took the ratio in intensity in the SAXS pattern between the direction of deformation and the direction orthogonal to deformation, A_asym_ (see Supplementary Fig. S1 and Eq. 3 for a definition). Shearing caused A_asym_ to increase from zero (isotropic state) to a maximum of nearly 0.5 at a shear strain of 250% (SI Fig. S1). As shown in Fig. 4b, *A*_*asym*_ returned back to zero when the sample was returned to 0% strain, over the entire range of shear strains. We conclude that shear-induced filament alignment is fully reversible, even up to strains close to the rupture point. Indeed, the fibrin networks are almost perfectly elastic due to covalent crosslinking by FXIIIa (see Supplementary Fig. S2), which should prevent fiber-fiber or monomer-monomer sliding^43,57,58^.

**Figure 4.**
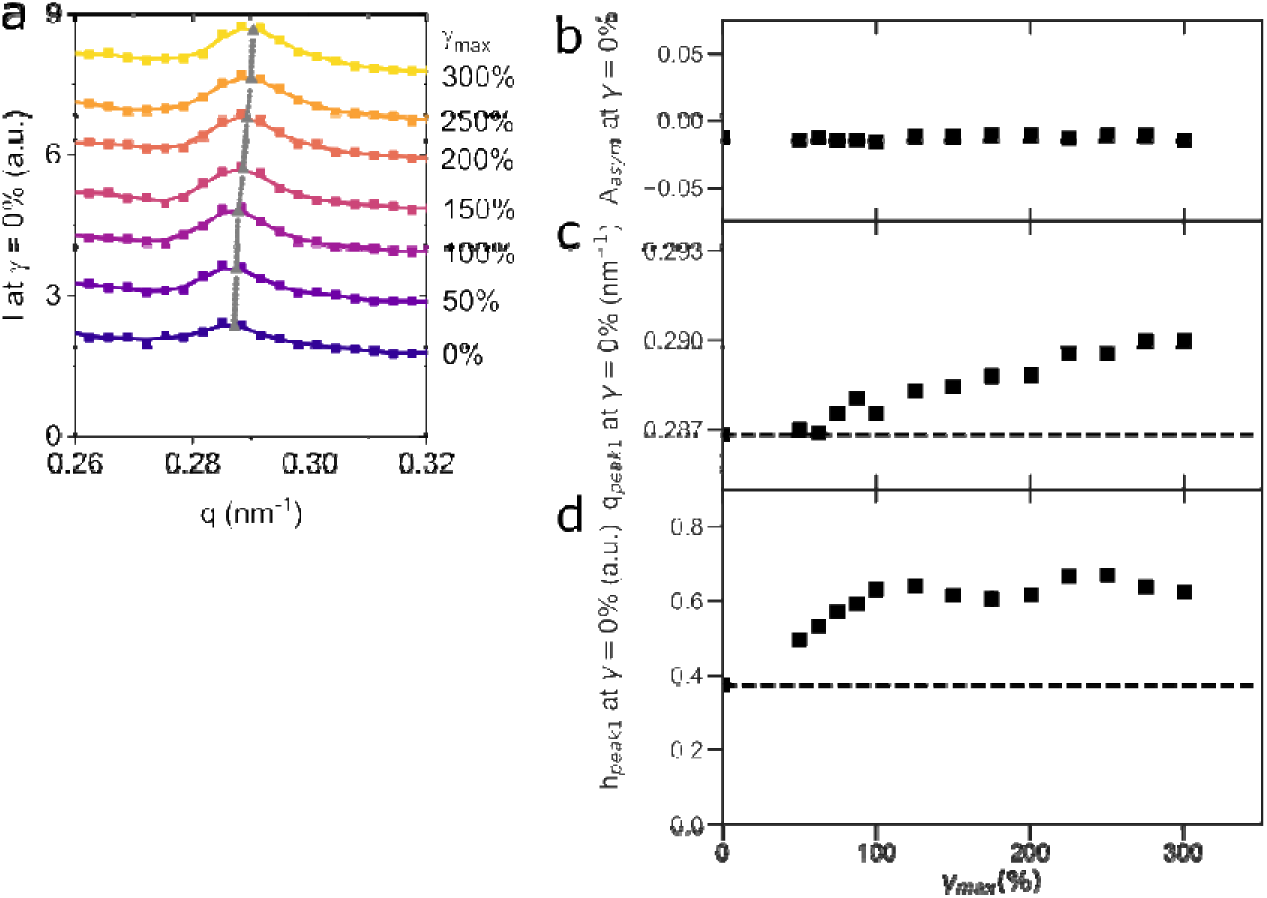
Reversibility of shear-induced changes in fiber and network structure. (a) The radially integrated intensities of the scattering patterns after return to zero strain after prior exposure to different levels of shear strains (legend on the right). The grey squares indicate the peak position *q*_*peak 1*_, obtained from fitting the radially integrated scattering curve. For clarity, the curves have been shifted along the y-axis. (b) Anisotropy in the scattering patterns calculated from the asymmetry factor (Eq. 3) measured upon return of fibrin networks to zero strain after being strained out to γ_max_. Fiber alignment is fully reversible upon removal of the strain. (c) The position of the first scattering peak measured upon return of fibrin networks to zero strain after being strained out to γ_max_. (d) The corresponding height of the first scattering peak. The two colours represent two independently measured data sets. The horizontal dashed lines indicate the value of the peak position and height of the origin, unstrained fibrin network. A_asym_, *q*_*peak 1*_, *h*_*peak 1*_ and error bars on *q*_*peak 1*_ and *h*_*peak 1*_ are the average between two independent measurements; the error bars are smaller than the black squares.

To analyse the reversibility of shear-induced changes in the fiber structure itself, we considered the position and height of the Bragg peak measuring the axial order (Fig. 4a). We compared the peak position and height for fibrin gels returned to zero strain after exposure to different levels of shear strain with the reference values that were originally measured for the gel in its original state (dashed horizontal lines in Fig. 4b,c). For strains up to 50%, both the position and the height of the peak did not appreciably change as the sample was sheared (Fig. 3a-b) and remained close to their original levels upon return to zero strain (Fig. 4c-d). By contrast, when the strain was increased beyond 50%, the Bragg peak shifted to lower *q*-values with increasing shear strain (Fig. 3a) whereas it shifted to higher q-values compared to the original state upon return to zero strain (Fig. 4c). For the most strained gel, the repeat length decreased by 0.5 nm with respect to the initial length. Moreover, the height of the peak increased with increasing previously experienced strain and was higher than the peak height measured in the original state (Fig. 4d). These observations indicate that, upon removal of the strain, the characteristic repeat length *shortens* with respect to the original length in the virgin network, and that the axial packing order is enhanced by stretching and relaxation. Apparently, the axial order of the fibrin protofibrils is imperfect in the virgin network and mechanical cycling anneals out the initial disorder.

## Discussion

The aim of our work was to elucidate how fibrin gels obtain their remarkable strain-stiffening behaviour, which spans a wide range of shear strains up to 250% (Fig. 3d). We therefore combined macroscopic measurements of the shear rheology of fibrin networks with *in situ* structural measurements by SAXS. We found that the networks exhibit multiple distinct mechanical regimes that are closely correlated with distinct changes in network structure on different length scales.

At low strains (γ < 25%), the fibrin networks underwent a transition from linear elasticity (γ < 2%) to a strain-stiffening regime, but we did not observe any concurrent change in fiber orientation. This observation is consistent with prior *in situ* neutron scattering measurements on sheared fibrin networks^59^ and supports the idea proposed in several studies^12,56,59^ that fibrin networks initially store entropic slack in thermal bending undulations of the fibers. Shearing the network pulls out the slack, so the shear modulus remains constant until the strain is large enough to pull the fibers taut.

At larger strains (γ > 25%), we observed progressive fiber alignment along the principal direction of strain, again in line with neutron scattering data^59^. We find that fiber reorientation follows the macroscopically applied shear strain, indicating that the network deforms affinely. This finding confirms earlier studies, which reached the same conclusion based on the close correspondence between the elastic modulus of fibrin networks and predictions of affine network models^12,45^. Although the fibers oriented along the principal direction of strain experience substantial tensile strain, we did not observe any changes in the Bragg peak reflecting the axially ordered structure of the fibers until the shear strain reached 100%. Beyond 100% strain, we finally did observe progressive changes in the position and height of the Bragg peak, but these changes were small. The peak position shifted to lower *q*-values with increasing strain, indicating molecular elongation. However, the maximum increase in molecular length reached just before network rupture was just 0.7%, even though the fiber length is increased by 83% at this point. Simultaneously with a shift in the peak position, the peak height decreased with increasing strain. Apparently, the packing structure of the fibers becomes more disordered upon elongation.

Both the minimal peak shift and the enhanced disorder that we observe for fibrin networks undergoing shear are entirely consistent with previous *in situ* SAXS measurements on fibrin networks undergoing uniaxial stretch^39,60^. The fact that the axial periodicity does not follow the macroscopic strain rules out a fiber elongation mechanism mediated by simultaneous stretching of the monomers, as it is for instance observed in collagen fibers^61^. Like fibrin fibers, collagen fibers are bundles of axially staggered, rod-shaped monomers. However, collagen monomers have a simpler tertiary structure than fibrin monomers, with the bulk of the molecule forming an ordered triple-helical rod. By contrast, fibrin monomers combine a folded trinodular portion having a length of 45 nm with two long and flexible regions that extend out from the two ends (390 residues each)^5,62,63^. The monomers form double-stranded protofibrils that laterally associate via the long and flexible αC-tethers^64–66^. Fibrin fibers therefore have multiple mechanisms available for elongation: they can either elongate by forced unfolding of the protofibrils, or by entropic stretching of the natively unfolded αC-connectors.

In principle both elongation mechanisms could explain the behaviour of the Bragg peak measured for fibrin networks under shear or uniaxial stretch. The first model assumes that the tensile force is supported by the γ-γ contacts of monomers within the protofibrils, and the monomers act as springs in series. The model furthermore assumes that the alpha-helical coiled-coil connectors and/or globular domains in the γ-nodules unfold^31,32,39,67^ (Fig. 5B). For this mechanism to hold, it is necessary to assume that the monomers unfold stochastically, so there remains a population of unfolded molecules that maintains the original 22.5 nm spacing. Since the Bragg peak is an ensemble-averaged quantity that averages over packing heterogeneities of a large ensemble of fibers, this model predicts that the network should exhibit a Bragg peak that has a constant position but gradually decreases in height as more and more monomers unfold, where it assumes that unfolded molecules no longer contribute to scattering. Full monomer unfolding can theoretically cause a 5-fold fiber elongation, assuming that each amino acid residue contributes a contour length of 0.38 nm in the unfolded state^31^. The γ-nodules (215 residues) can increase in length from 5 to 80 nm and the coiled coils can increase in length from 17 to 41 nm. Single-molecule measurements and simulations showed that in the absence of αC-αC interactions, both domains indeed unfold under axial loading^10,31,32,68^.

**Figure 5.**
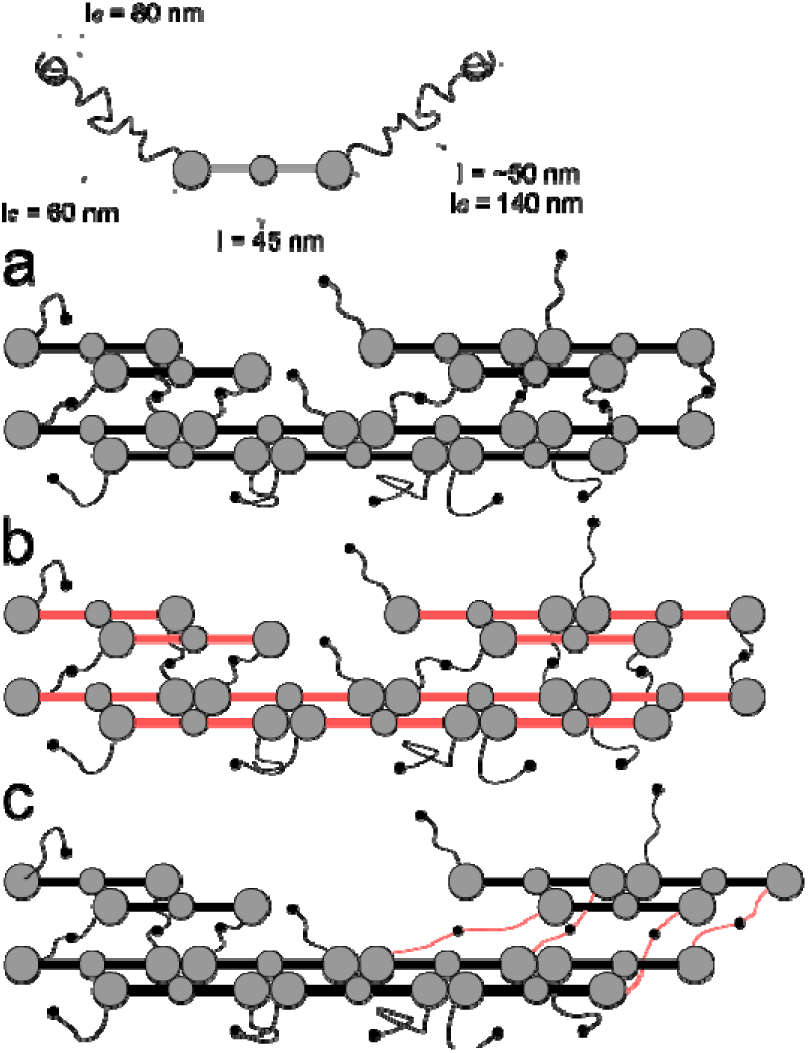
Schematic representation of two different mechanisms of fibrin fiber extension. (a) The schematic shows a fibrin monomer with length l of the trinodular core and the effective length l and contour length *l*_*c*_ of the entire αC-region (αC-connector and the αC-domain). Shown below are sections of two connected protofibrils at rest, with an axial packing periodicity of 22.5 nm that corresponds to half the monomer length. The packing defect in the top row arises due to the finite length of the protofibrils within fibrin fibers. (b, c) A schematic representation of two elongation mechanisms for fibrin fibers, (b) through forced unfolding of the monomer, and (c) through entropic stretching of the natively unfolded αC-connectors. The length of the αC-region is obtained from Ref.^95^, the contour length of the αC-region, domain and connector are obtained from Ref.^63^, and the length of the fibrinogen monomer from Ref.^47,96^.

The second mechanism for fiber elongation instead assumes that tensile strain is borne by the flexible αC-regions in between protofibrils^37,69^ (Fig. 5C). The rationale for this mechanism is that the αC-region consists of a natively unfolded αC-connector of 170 amino acids that can be modelled as a wormlike chain with a persistence length of 0.6 nm, and a folded αC-domain of 218 amino acids^11,63^. The αC-region as a whole can accommodate large elongations because it is only 45-50 nm long at rest^70^ and can be stretched to 148 nm. If the fiber elongates through stretching of the αC-tethers, we expect no shift of the Bragg peak until the strain reaches a point where the αC-tethers are pulled taut and the protofibrils start to feel the strain. We furthermore expect Bragg peak broadening due to increased axial disorder upon fiber elongation, because the protofibrils lose registration upon sliding.

To distinguish between the two mechanisms, we consider the rheology data, which provides insight into the tensile force experienced by the fibers. At the onset strain of 100% where we start to observe a shift in the Bragg peak position, we measured a shear stress of 1330 Pa. Assuming that the stress is distributed uniformly over the network and knowing that the fibers consist on average of *N*_*p*_ = 288 double-stranded protofibrils and the mesh size is 1.35 µm (as inferred from turbidimetry), we estimate a tensile force per monomer of 8.4 pN at this point. This force is much lower than the minimum unfolding force of 50 pN for the γ-nodules and coiled coils observed in single-molecule AFM experiments^10,31^ and in simulations^31–33,71^, making forced unfolding of protofibrils unlikely. Instead, it is much more likely that the natively unfolded αC-tethers initially accommodate the applied strain. Since the connector region is unstructured, it should stretch as a random coil polymer, without a critical force and with a continuous extension up to a force of 25 pN^37^. Several lines of evidence in the literature support this model. First, force-extension measurements on single fibrin fibers prepared from fibrinogen from different animal sources showed that the maximum fiber extensibility increased linearly with the length of the αC-region^36,72^. Second, single fiber force-extension curves have been shown to be linear up to strains of 100%, consistent with progressive stretching of the αC-tethers and inconsistent with unfolding^73^. Third, the Young’s modulus of fibrin fibers is on the order of 1-10 MPa, consistent with the low-strain elasticity being dominated by the flexible αC-tethers rather than the rigid (GPa) coiled-coils^35^. Fourth, single fibers stretched out to 100% exhibit fast recoil within <4 ms and this recoil is repeatable over multiple stretch cycles, as expected from the wormlike chain behaviour of the αC-tethers^37^. Molecular modelling suggests that refolding of unfolded γ-nodules is too slow to account for such fast recoil^37^.

Our data furthermore suggest that, once the shear strain exceeds 100%, the fibers elongate by a combination of αC-tether elongation and protofibril stretching since we observe a progressive, albeit small, elongation of the axial periodicity. Concurrently, we observe a steeper rise of the elastic modulus at the largest strains, signifying a nonlinearity in the force-extension response of the fibrin fibers. The source of this nonlinearity could be either (partial) unfolding of the γ-nodules, which would cause the formation of unstructured peptide stretches that stiffen entropically, or unfolding of the coiled-coils, which convert from an α-helical to a more rigid β-sheet structure^34^. We observe a maximum molecular elongation of 0.3 nm just before the network ruptures. This is comparable in order of magnitude to the shift reported in a recent SAXS study of pre-aligned plasma clots under uniaxial strain, where the shift was 5% when the tensile strain reached 40% (at which point the peak disappeared)^60^. Several studies showed direct evidence of forced unfolding of fibrin monomers within fibrin gels subject to large shear or tensile loads, using wide-angle X-ray scattering^74^ or vibrational spectroscopy^41,42^. Indeed, due to axial and transverse crosslinking of the protofibrils, it is likely that stretching of the αC-regions is somehow coupled to protofibril elongation.

We expect that the strain partitioning between the protofibrils and the αC-regions will depend on the exact connectivity of the fibers, which is a function of the length of the protofibrils. Short protofibril lengths will tend to favour αC-tether stretching, whereas long protofibril lengths will tend to favour protofibril stretching. It is difficult to measure the length of the protofibrils directly. Several studies on the kinetics of fibrin assembly suggest that protofibrils are relatively short (up to 20 monomers) before they start to laterally associate under standard polymerization conditions^75–77^. However, the assembly kinetics of fibrin are known to be strongly dependent on the fibrinogen and thrombin concentrations, the temperature, and the pH and ionic conditions^75,78^. It is therefore possible that fibrin fibers assembled under different conditions will not have the same connectivity and will differ in the contributions of αC-tether and protofibril stretching. The strain distribution is also likely to depend on the levels of crosslinking (SI Fig. 2). FXIIIa creates γ-γ-crosslinks along the longitudinal direction of the monomer, which may channel stress through the D and E nodules and the coiled coils^11^. FXIIIa furthermore creates multimeric crosslinks between the multiple donor residues in the flexible αC-connector and multiple acceptor residues in the globular αC-domain. Enhanced α-α crosslinking will constrain the conformation of the αC-regions and limit their extensibility^11^.

An intriguing finding of our work is that the molecular packing distance decreases upon the return of the fibrin network from a highly deformed state to zero strain. This observation is reminiscent of a recent SAXS study that similarly reported a permanent small decrease in axial periodicity on relaxation after prior network stretching. However, contrary to that study, we observe that the packing order is enhanced compared to the virgin state, whereas the earlier study found that the packing order was reduced. We propose that enhanced packing order after shear is consistent with the model of fiber elongation we proposed above: stretching of the αC-tethers will cause sliding among protofibrils, causing them to lose registration. The lateral packing structure of protofibrils is known to be rather disordered^27,48,79–83^ and strongly influenced by the assembly conditions^29,84–88^. Stretching and subsequent relaxation may therefore allow the protofibrils to find a more ordered packing configuration and anneal out initial imperfections, similar to strain annealing in nanocrystalline metallic materials^89^.

## Conclusion

We performed X-ray measurements coupled with shear rheology to reveal the structural mechanisms that mediate the nonlinear elastic response and large resilience of fibrin networks. We showed that increasing levels of shear strain induce distinct changes in elasticity that coincide with a sequence of structural responses. There were no observable changes in the fiber or network structure up to strains of around 25%, consistent with entropic elasticity. Above 25% strain the fibers progressively aligned along the principal direction of strain, causing network stiffening. We found that fiber reorientation closely followed the expected alignment for an affinely deforming network, showing that the strain was uniformly distributed down to length scales of at least 50 nm. Once the strain reached values above 100%, we observed a progressive increase of the axial packing distance, albeit much smaller than the total fiber strain. Surprisingly, when the strain was brought back to zero, the fiber orientations returned back to the original state, whereas the axial packing distance slightly shrank compared to the original state, and the axial packing order increased. To explain these findings, we propose a model where fibrin fibers are a composite structure of rigid protofibrils connected by flexible tethers into relatively disordered arrangement that can be remodelled by mechanical stretching. Interestingly, other biological elastomers use either unfolding of ordered^7,8^ or disordered domains^9^ to accommodate large deformations, while fibrin uses a combination of both pathways. Our findings provide a mechanistic framework to understand the strain-dependent mechanical response of blood clots, which is a prerequisite to understand the molecular basis of blood clotting disorders. Moreover, our findings identify hierarchical structuring as a powerful design principle to make resilient materials.

## Acknowledgements

We gratefully acknowledge Paul Kouwer (Radboud University Nijmegen, the Netherlands) for kindly lending us his TA HR2-rheometer during beamtime BM26-02-797. We also acknowledge the beamline staff, in particular Daniel Hermida Merino, Giuseppe Portale and Wim Bras, of BM26 at the ESRF for their help with the experiments and data analysis. We thank Bela Mulder (AMOLF, Amsterdam, the Netherlands) for discussions on the calculation of the nematic order parameter, and Karin Jansen and Lucia Baldauf (AMOLF, Amsterdam, the Netherlands) for contributing to the measurements and discussion on the analysis of SAXS-data. We thank Martin van Hecke for lending us the confocal rheometer head, Marnix Verweij, Jan Bonne Aans and Dirk-Jan Spaanderman for help with the construction of the confocal rheometer and for designing the SAXS-adaptation for the Anton Paar rheometer. This work was financially supported by NWO, through a FOM Program grant (nr 143) and DUBBLE-beamtime proposals BM26-02-797, BM26-02-857, and SC-4789. The work of F.B. and G.H.K. is part of the Industrial Partnership Programme Hybrid Soft Materials that is carried out under an agreement between Unilever Research and Development B.V. and the Netherlands Organisation for Scientific Research (NWO).

## Materials and methods

### Fibrin network assembly

Human plasma fibrinogen (Plasminogen, von Willebrand Factor and Fibronectin depleted) and human α-thrombin were obtained in lyophilized form from Enzyme Research Laboratories (Swansea, United Kingdom). All chemicals were obtained from Sigma Aldrich (Zwijndrecht, The Netherlands). Fibrinogen (lyophilized in 20 mM sodium citrate-HCl buffer at pH 7.4) was dissolved in water at 37°C for 15 min to its original concentration (ca. 13 mg/ml) and extensively dialysed against fibrin buffer containing 20 mM HEPES and 150 mM NaCl at a pH of 7.4 in order to remove citrate, which complexes with Ca^2+^ ions that are required for FXIIIa and thrombin activity. After dialysis, the fibrinogen solution was aliquoted, snap-frozen in liquid nitrogen and stored at −80°C. The stock concentration was around 10 mg/ml, as determined by UV-VIS spectrophotometry at 280 nm using a Nanodrop 2000 spectrophotometer (Thermo Scientific) and assuming an extinction coefficient of 16.0 mg/(ml.cm)^90^. Thrombin (lyophilized in 50 mM sodium citrate and 0.2 M NaCl) was reconstituted in water to its original concentration (approximately 10,000 U/ml), and quickly aliquoted, snap-frozen in liquid nitrogen and stored at −80°C.

Prior to use, fibrinogen and thrombin aliquots were thawed at 37°C and kept for at most 1 day at room temperature and on ice, respectively. Fibrinogen was mixed with 500 mM CaCl_2_ and assembly buffer at room temperature to reach a final concentration of 4 mg/ml in a final assembly buffer containing 20 mM HEPES, 150 mM NaCl and 5 mM CaCl_2_. Fibrin network assembly was initiated by adding 0.5 U/ml of thrombin from a 20 U/ml thrombin stock. After mixing, the mixture was quickly transferred to the rheometer, to allow *in situ* polymerization at 37°C.

### Turbidimetry measurements

To characterize the fiber diameter and mass-length ratio, we performed turbidimetry measurements using a Lambda 35 spectrophotometer (Perkin Elmer, Groningen, The Netherlands). We used quartz cuvettes suitable for fluorescence measurements (Hellma, Kruibeke, Belgium) with a 10.0 mm or 2 mm path length. Prior to measurements, a baseline correction was performed by placing a cuvette containing only the assembly buffer in both the measurement and reference position. Measurements were performed at room temperature. Wavelength scans were taken from 450 nm to 900 nm at 1 nm intervals at a scanning rate of 480 nm/s and a slit width of 2 nm. Data were analysed using a model for isotropic networks of rigid fibers published in Ref.^91^, including a correction for wavelength dispersion of the refractive index of the solvent n and the differential refractive index^91^.

### Rheology

Rheological experiments were performed using a Kinexus rheometer (Kinexus pro+, Malvern, United Kingdom) equipped with a 40 mm stainless steel cone-plate geometry with a 1° angle. During polymerization, the temperature was kept constant at 37°C, and solvent evaporation was prevented by adding a layer of mineral oil (M3516, Sigma Aldrich) on the liquid-air interface. We checked that the oil layer did not influence the mechanical behaviour of the networks by comparing experiments in the absence and presence of an oil layer. We found insignificant differences between the mechanical response of samples with and without oil, falling within the variability between repeat measurements. Full-length fibrinogen was polymerized *in situ* without surface treatment. During polymerization, we continuously measured the shear modulus by applying a small oscillatory shear strain with an amplitude of 0.5% and a frequency of 0.5 Hz. Polymerization was complete after 2 hours, as indicated by a time-independent shear modulus. We measured the mechanical response to increasing levels of stress by applying a constant shear stress, superimposed with an oscillatory shear stress with an amplitude of 0.1 times the constant shear stress and a frequency of 0.5 Hz. We then increased both the constant and oscillatory stress level, to maintain their 10-to-1 ratio, until the sample broke. As the applied oscillatory stress is small compared to the total stress on the network, we calculate the differential modulus *K* = d*σ*/d*γ* ≈ Δ*σ*/Δ*γ*, with Δ*σ* the applied amplitude of the oscillatory stress, and Δγ the measured amplitude of the oscillatory strain.

### SAXS measurements

Small Angle X-ray Scattering (SAXS) experiments were performed at the DUBBLE BM26B beamline at the ESRF (European Synchrotron Radiation Facility) in Grenoble, France. The energy of the x-rays was 12 keV, corresponding to a wavelength of 1.0*10^−10^m. The sample-to-detector distance was adjustable; for our experiment we chose a distance of 3 m to optimally access the relevant *q*-range possible with this setup. The specified beam dimensions were 300 x 300 µm.

A Pilatus 1M detector^92^ (169 mm x 179 mm active area) was used to collect SAXS images. A beamstop was placed in between the sample and detector to block the unscattered X-ray beam and thus prevent overexposure and damage of the detector. Both the beamstop and its holder can be seen in Fig. 2a in the Supporting Information in a so-called mask image: a binary image where all pixels that are not suitable for analysis are indicated in black. Determination of the center position of the scattering images, as well as calibration of the wavevectors, was done with a sample of AgBe (silver behenate) powder (Fig. S3b in the Supporting Information).

Measurements of fibrin gels under deformation were performed by polymerizing the fibrin sample in a home-built polycarbonate Couette cell attached to an Anton Paar MCR501 (Anton Paar, Graz, Austria) rheometer. The rheometer’s Peltier cell maintained a constant temperature of 37°C during assembly. Per measurement, 2 mL of sample was used. The Couette is designed such that the outer wall is stationary, and the inner cylinder is attached to the rheometer spindle and rotates. During rheology experiments, a background image of the assembly buffer was taken for 60 s prior to every measurement. For the fibrin samples, the acquisition time was 45 s per strain step. No visual damage from the high-energy radiation to the sample was observed during the SAXS measurements.

### SAXS data analysis: background subtraction and integration

We wrote a customized Python program to analyze the scattering images, based on the following procedure. From the AgBe calibration image (SI-Fig. 3b), we obtained the center position of the beam on the detector (x_0_, y_0_). For every pixel in the scattering image, we calculated the distance *d* = [(*x* − *x*_0_)^2^+(*y*-*y*_0_)^2^]^0.5^ and the angle θ = atan[(*y* − *y*_0_)/(*x* − *x*_0_)] to the center. Next, the bands in between the detector panels and the beamstop were removed by the application of a binary mask and the background was subtracted (SI-Fig. 4). Since the scattering signal from the fibrin network is weak, careful background subtraction is essential (see Fig. S4 in the Supporting Information). To convert distances in pixels to wavenumbers (q in units of nm^−1^), we use the following formula:

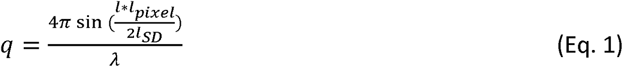

where *l* is a length in pixels, *l*_pixel_ = 172 µm the size of an individual pixel, *l*_SD_ the sample-to-detector distance (around 3 m) and *λ* the wavelength of the X-rays.

Radial integration of the scattering intensity was performed by taking the average value of all pixels that fell in a distance bin size with a width of 1 pixel. Azimuthal integration was performed by averaging all pixels within a specified *q*-range corresponding to 50 nm to 102 nm, with a bin size with a width of θ = 2°. The lower limit was set by the limitation that we do not want to take the Bragg peak originating from the molecular packing order into account, while the upper limit was set by the lowest q-value we can access. We verified that using this *q*-range we did not include the panel boundaries in the azimuthal integration. A slice of θ = 40° was left out at the top to eliminate the holder of the beamstop.

### SAXS data analysis: Bragg scattering peak

To determine the position of the Bragg-peak resulting from the periodic axial packing structure of fibrin fibers, we subtracted the slope of the scattered intensity that is convoluted with the Bragg peak. In detail, we low-pass filtered the data by smoothening with a Gaussian filter with a small width (*σ* = 1 pixel, corresponding to approximately q = 0.0034 nm^−1^ at this *q*-range). We then applied deconvolution with a kernel of the derivative of a Gaussian with a large width to remove high-frequency noise (without smoothing the Bragg peak) and obtain the local slope at the peak position. As a result, we obtained the Bragg peak with a flat baseline (SI-Fig. 5). We fitted this peak to an analytical function to quantify its shape. In literature several functions are used to describe a Bragg peak, viz. Gaussian, Lorentzian, Voight, pseudo-Voight, or Pearson VII^93^. However, because of low signals in our experiment, we cannot distinguish the exact shape, hence we simply used a Gaussian function to characterize the peak (in line with an earlier SAXS study of fibrin^39^). The Gaussian function has the form:

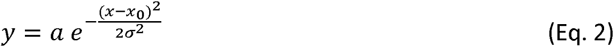

with *a* the peak height, *x*_0_ the peak center, *σ* the standard deviation and *b* the offset. In the main text, we report values for *x*_0_ as a measure of the peak position and a as a measure of the degree of crystalline order.

### SAXS data analysis: quantification of fiber alignment

To quantify shear-induced fiber alignment along the principal strain direction, we consider the geometry of the shear deformation relative to the X-ray beam (SI-Fig. 6). We apply shear by moving the inner cylinder of a Couette cell while keeping the outer cylinder stationary. Shearing sets up a strain field in the *ϕ* -plane, set by the gap size between the two cylinder walls. As a result, the projected length of the filaments will change both on the *ϕ* -plane and on the θ-plane, defined by the surfaces of the cylinders. The X-ray beam passes perpendicularly through the surfaces of the Couette cell, so the scattering image captures the projection on the θ-plane.

To quantify the anisotropy from the scattering patterns we used two methods. The first method is a model-independent approach, where we quantify anistropic simply from the ratio between the intensities of a horizontal quadrant and a vertical quadrant:

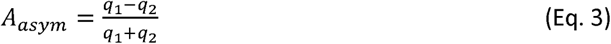

where *q*_1_ and *q*_2_ are the total scattering intensities in the orthogonal quadrants 1 and 2, respectively (see SI-Fig. 1). This parameter is rather sensitive to the background subtraction procedure, but it has the benefit that it can be applied even to samples at low strain that lack clearly distinguishable peaks in the orientational distribution. In the second method, we obtained the nematic order parameter from fitting an Orientational Distribution Function (ODF) to the azimuthally integrated scattered intensity as a function of *ϕ*, following an d approach originally developed to quantify nematic ordering of biopolymers from SAXS measurements^54,55^. This method has the advantage of permitting a quantitative comparison of shear-induced order with literature, but the downside is a large uncertainty in the fit to samples at low strain that lack clearly distinguishable peaks in the orientational distribution. The full derivation is given in Supplementary Document 1.

### Simulations of affine deformations on randomly oriented rods

To benchmark the development of shear-induced alignment in fibrin networks, we calculated the *affine* effect of shear deformation on an initially isotropic network using simulations of initially random, three-dimensional ensembles of rigid, elastic rods. Each rod had a fixed initial length of r = 1, and was further characterized by two random angles θ and φ. We generated a sufficiently large number of rods (100,000) to obtain a statistically significant ensemble average, and applied strain in small steps of 0.01 using a deformation matrix for simple shear in 3 dimensions:

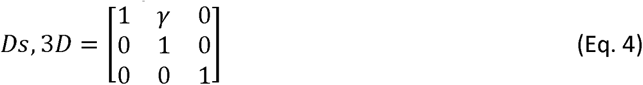

After a deformation matrix is applied to each filament, we re-calculate its length and orientation. To quantify the (average) orientation in the deformed network, we calculate the nematic order parameter *S*^94^, starting from the tensor order parameter *Q*:

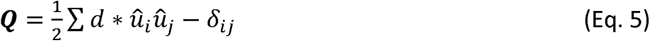

with *δ*_*i,j*_ the Kronecker delta function that is 1 when *i* = *j* and 0 otherwise. Based on a given set of rod angles θ and *ϕ*, the tensor order parameter reads:

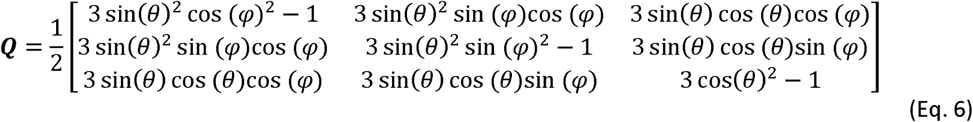

By quadratically summing all terms and taking the square root, we obtain the nematic order parameter, invariant to the orientation of the strain:

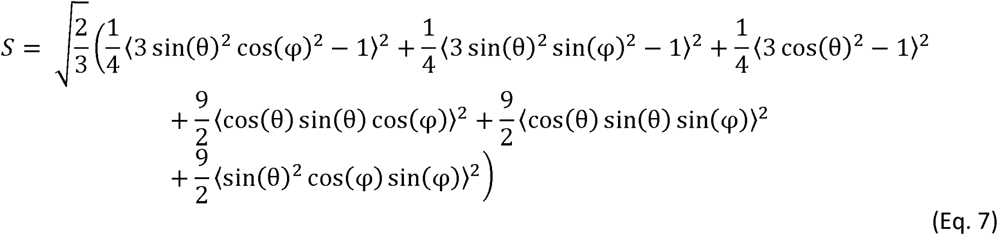

Note that *S* is normalized, such that it equals zero for an isotropic system and one for a perfectly aligned system.

From the simulations, we can also compute the asymmetry factor corresponding to the geometry of our experiment, by considering the projections of the rods on the θ and *ϕ* plane. In SI-Fig. 7a the probability function is shown for the *φ* -coordinate for an ensemble of filaments under shear deformation. At zero strain, this function is flat. As the shear angle changes with increasing strain, the peak position of the probability function shifts for increasing strain levels, and increases in intensity. The probability function is periodic with a shift of π, as a filament with length (*x*_*i*_, *y*_*i*_) is identical to a filament with length (-*x*_*i*_, -*y*_*i*_). In SI-Fig. 7b the probability function is shown for the ϕ-coordinate. The probability distribution at zero strain behaves as a cosine function, with a peak developing at ϕ = π/2 as strain increases, indicating alignment towards the *φ* -plane.

From the simulations, we can also obtain the distribution of fiber strains. In SI-Fig. 7c we display the probability function for the length of the filaments. At zero strain, this is a single spike at *l* = 1, but as strain increases, the distribution broadens. As the strain orientation changes during shear, some individual filaments experience a higher strain than the global shear strain. There is not only a population of filaments that is extended, but also a population of filaments that is compressed as the filaments are oriented orthogonal to the direction of strain. As shown in the inset, the fraction of the filament population that is under compression, extension up to 100% and extension of more than 100% is shown. Even for 300% applied shear, approximately 20% of the filaments are still under compression.

### Confocal rheology

In order to visualize the changes in fibrin network architecture under shear directly, we combined shear rheology with confocal imaging using a home-built setup consisting of a rheometer head (Anton Paar DSR 301) on top of an inverted microscope (Leica DM IRB) equipped with a Yokogawa CSU-22 spinning disk, a 100x oil objective, and a Hamamatsu EM-CCD C9100 camera. The rheometer head could be manually lifted and lowered using a micrometer screw. The microscope objective was situated underneath the glass coverslip, such that the center of the objective was located halfway the center and the edge of the rotating top plate. A fibrin gel spiked with 10 mole% of fluorescently labelled (Alexa-488) monomers was polymerized *in situ*, in between a glass coverslip and a 20 mm diameter stainless-steel plate attached to the rheometer. A layer of low-viscosity mineral oil (M3516, Sigma Aldrich) was applied at the sample-air interface to prevent solvent evaporation. Network polymerization was monitored by small amplitude oscillatory shear deformations with a strain amplitude of 0.5% and frequency of 0.5 Hz. At room temperature (22°C), polymerization was complete after 2 hours.

## Notes

#### Summary of Updates

Corrected author information (Koenderink)

