## Supplementary Figures for "Revealing the molecular origins of fibrin’s elastomeric properties by in situ X-ray scattering"

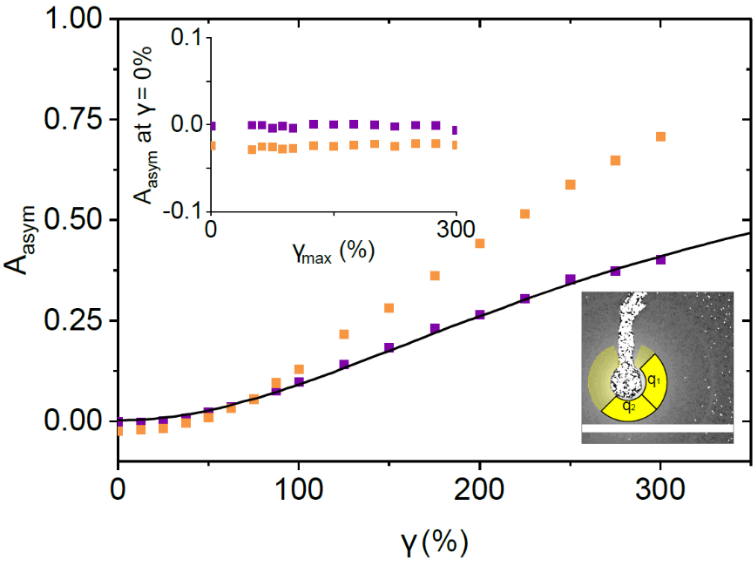


**Supplementary Figure 1. Asymmetry factor $A_{asym}$ as a function of applied shear strain γ for a 4 mg/ml fibrin gel.** Symbols display data for two independently prepared samples. The black line shows the model prediction in case of an affinely deforming ensemble of rods. In the main text we fitted an Orientational Distribution Function to the azimuthally integrated scattered intensity to obtain the nematic order parameter (Fig. 3e). Here, we report the ratio in intensity in the SAXS pattern between the direction of deformation and the direction orthogonal to deformation (Eq. 3). The advantage of the anisotropy factor $A_{asym}$ is that it is model-independent and applicable at low applied strain when there is negligible anisotropy, while the downside is that it is more sensitive to noise than the nematic order parameter and not directly comparable with literature. The top left inset shows $A_{asym}$ when the sample is returned to 0% strain, as a function of the maximum applied strain $\gamma_{max}$. The bottom-right inset shows the definition of q1 and q2, as used for radial integration.

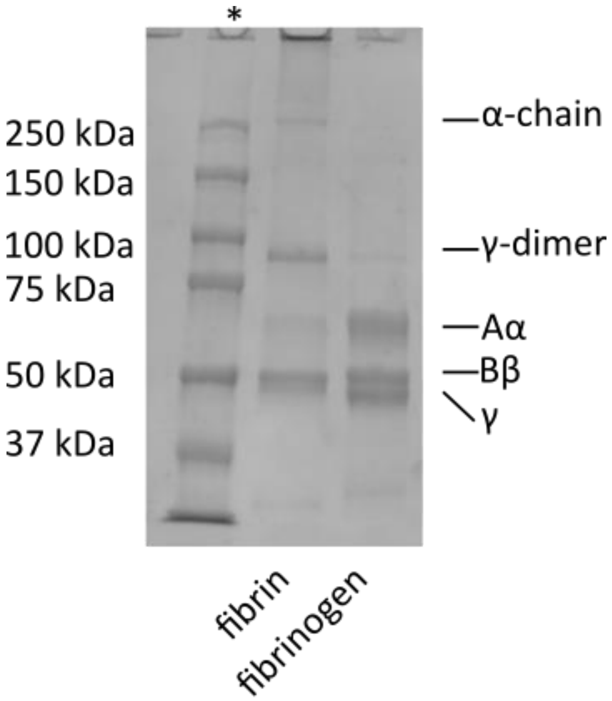


**Supplementary Figure 2. SDS-PAGE gel of fibrinogen and of the fibrin gel obtained upon reaction with thrombin.** We checked the purity and the degree of crosslinking of our fibrinogen and fibrin gels on a 7.5% SDS-gel, using 2.5 µg of protein per lane, staining with InstantBlue (Gentaur, Eersel, The Netherlands). The Aα, Bβ and γ-chains are indicated. Upon polymerization, the molecular weight of two polypeptide chains decreases due to the removal of fibrinopeptide A from the Aα-chain and fibrinopeptide B from the Bβ-chain. Furthermore, a new band appears around 100 kDa due to the formation of γ-dimers, indicative of crosslinking by FXIII that is co-purified with fibrinogen. Also high molecular weight bands appear due to the formation of α-chain crosslinks. Molecular masses were calibrated by running a lane with Bio-Rad Kaleidoscope Standards 161-0375 in the lane marked with an asterisk.

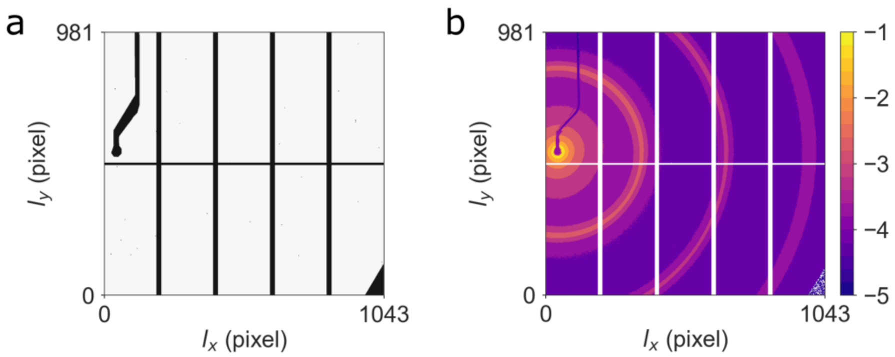


**Supplementary Figure 3. Control SAXS images used for data analysis.** (a) The mask used to filter out the beamstop and the bands in between the panels on the detector. Black pixels are removed from the scattering data during analysis. (b) AgBe sample used for calibration of the wavevector and determination of the center position. The first three scattering orders are clearly visible, and are used all three to determine the position of the X-ray beam and to calibrate the *q*-range. Data is shown on a logarithmic scale, in arbitrary units (see color bar).

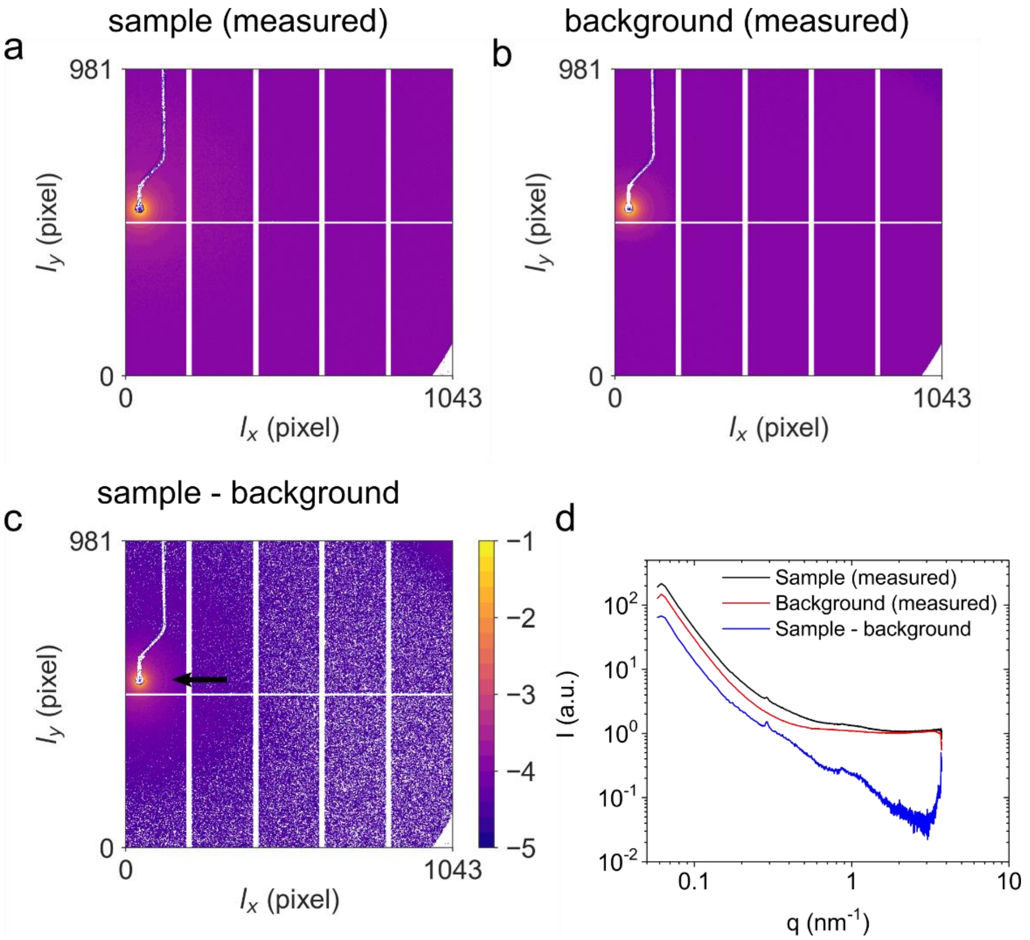


**Supplementary Figure 4. Procedure for background subtraction of SAXS scattering patterns.** (a) Scattering pattern recorded for a 4 mg/ml fibrin sample. (b) Scattering pattern recorded for the buffer background. (c) Background-subtracted image of the fibrin sample. Note that after background subtraction, the first order scattering ring is (dimly) visible around the beamstop, as indicated by the black arrow. The spacings between the panels and the beamstop are visible in the images, but are removed during data processing. Data is shown on a logarithmic scale, in arbitrary units (see color bar). (d) Radially integrated scattering intensity as a function of wavevector, showing the measured intensity for the 4 mg/ml fibrin sample in black, the background in red, and the background-subtracted intensity in blue. Note that the background-subtracted intensity of the fibrin network is approximately a factor 3 lower than the total measured intensity. At high q-values, the scattering intensity from the sample becomes nearly identical to that of the background, so small fluctuations (e.g. read-out noise) can cause the background-subtracted sample to show negative intensities in individual pixels.

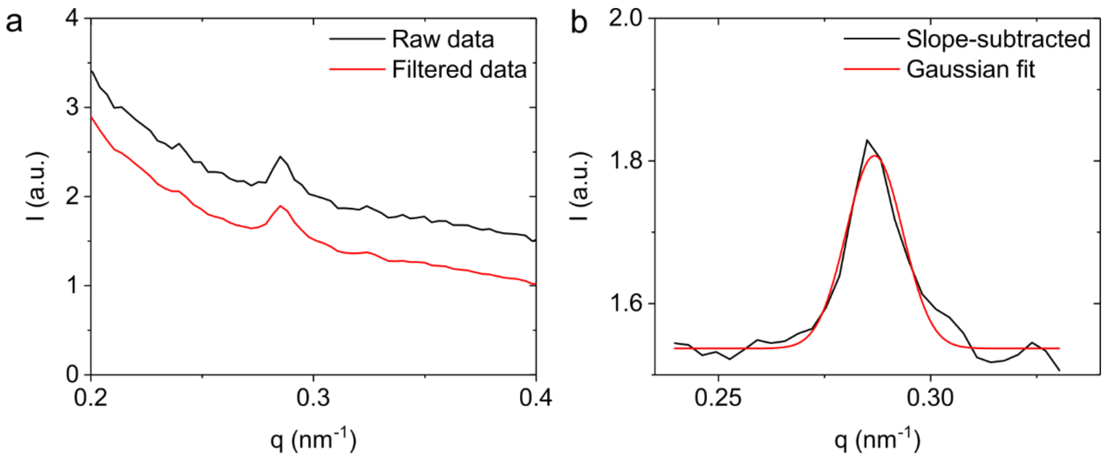


**Supplementary Figure 5. Peak-fitting procedure to analyze the position and height of the Bragg scattering peak.** (A) Example data (black line) measured for a 4 mg/ml fibrin gel and the data are filtered with a Gaussian with a small width (red line). For clarity, the curves have been shifted along the vertical axis. (b) Subtraction of the filtered data with the slope of the scattering curve, which was obtained by convoluting *q* with the derivative of a Gaussian with a large width (black line), and a fit with a Gaussian function (red line, Eq. 2). The fitting range runs from 24.0 $nm^{-1}$ to 32.4 $nm^{-1}$ and is chosen such that the Bragg scattering peak is centered in the middle of the fitting range, with one peak width extra on either side.

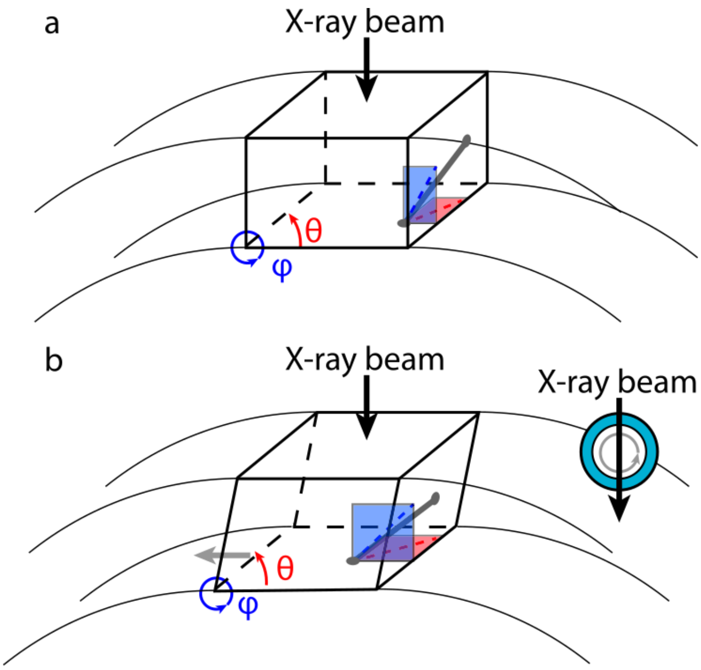


**Supplementary Figure 6. Geometry of the combined rheology/SAXS experiment.** (a) Schematic representation of the geometry of the experiment, where the X-ray beam hits the concentric inner and outer walls of the Couette cell. A single filament (dark grey) is sketched within an undeformed network (not shown). Its projection on the φ- and θ-plane is shown in blue and red, respectively. (b) As a shear strain (grey horizontal arrow) is applied in the φ-plane by rotating the inner cylinder, the filament stretches and aligns in the direction of principal strain. This reorientation also affects the projected length (red dashed line) on the θ-plane, which is the plane that is imaged through X-ray scattering. The inset shows a zoomed-out, top view of the Couette cell.

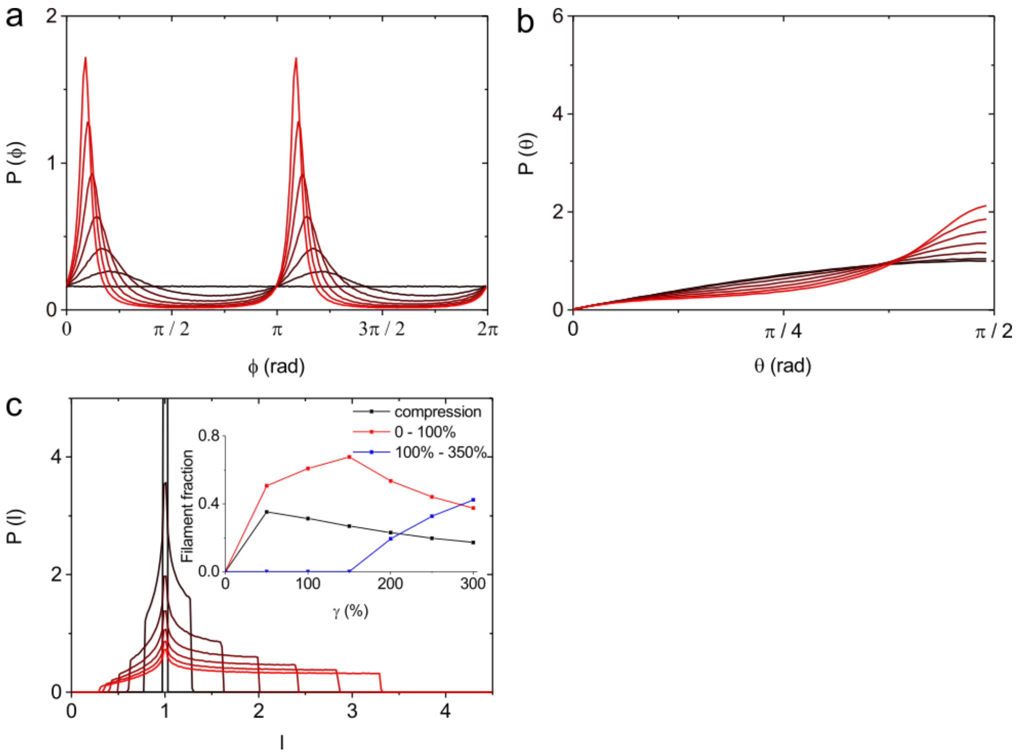


**Supplementary Figure 7. Predictions of filament alignment from affine simulations.** (a) Normalized probability function *P* for the φ-projection of a 3D ensemble of filaments under shear, with increasing applied deformation γ. (b) Corresponding probability function in the θ-direction. (c) Probability function for the filament length *l*. The inset shows the fractional population of filaments under compression, extension up to 100% and extension higher than 100%, as a function of the applied network strain γ. Orientations of 1e5 filaments were averaged to obtain graphs (a) and (b), 3e6 filaments were used to obtain graph (c).
