## Supplementary Document for "Revealing the molecular origins of fibrin’s elastomeric properties by in situ X-ray scattering"

### Supplementary Information: derivation of the nematic order parameter

The nematic order parameter $S$ was calculated using an approach originally developed to quantify liquid crystal formation of semiflexible fd-virus rods [1] and actin filaments [2]. We start with an expression for the Orientational Distribution Function (ODF) [2]:

$$\Phi(\theta) = Ae^{m\cos(\theta)^2} \tag{1}$$

where $A$ is a normalization constant. We determine the normalization constant $A$ by integrating $\Phi(\theta)$ over a solid angle $d\Omega$

$$\begin{aligned}
\Phi(\theta) &= \int_0^{2\pi}\int_0^{\pi} Ae^{m\cos(\theta)^2}\sin(\theta)d\phi d\theta && (2)\\
&= 2\pi A\int_0^{\pi} e^{m\cos(\theta)^2}\sin(\theta)d\theta && (3)\\
&= \frac{2\pi A}{\sqrt{m}}\int_0^{\pi} e^{(\sqrt{m}\cos(\theta))^2}(-1)d\sqrt{m}\cos(\theta) && (4)\\
&= \frac{2\pi A}{\sqrt{m}}\int_{\sqrt{m}\cos(0)}^{\sqrt{m}\cos(\pi)} e^{x^2}(-1)dx && (5)\\
&= \frac{2\pi^{3/2}A}{\sqrt{m}}\frac{1}{\sqrt{\pi}}\int_{\sqrt{m}}^{-\sqrt{m}} e^{x^2}(-1)dx && (6)\\
&= \frac{2\pi^{3/2}A}{\sqrt{m}}\frac{1}{\sqrt{\pi}}\int_{-\sqrt{m}}^{\sqrt{m}} e^{x^2}dx && (7)\\
&= \frac{2\pi^{3/2}A}{\sqrt{m}}\text{erfi}(\sqrt{m}) && (8)\\
&= 1 && (9)
\end{aligned}$$

using the imaginary error function $\text{erfi}(x) = \frac{1}{\sqrt{\pi}}\int_x^x e^{-t^2}dt$, $\frac{d\cos(\theta)}{d\theta} = -\sin(\theta)$, and changing variables $d\sqrt{m}\cos\theta = dx$, changing the integration boundaries accordingly. Hence,

$$A = \frac{\sqrt{m}}{2\pi^{3/2}\text{erfi}(\sqrt{m})} \tag{10}$$

so that

$$\Phi(\theta) = \frac{\sqrt{m}\,e^{m\cos(\theta)^2}}{2\pi^{3/2}\text{erfi}(\sqrt{m})} \tag{11}$$

We then add the fit parameters $a_1$, $a_2$ and $a_3$

$$\Phi(\theta) = a_1\frac{\sqrt{m}\,e^{m\cos(\theta-a_3)^2}}{2\pi^{3/2}\text{erfi}(\sqrt{m})} + a_2 \tag{12}$$

We then fit Equation 12 to the azimuthally integrated scattering intensity (see Fig. 3a), where $a_1$ and $m$ are free parameters, $a_2$ is set to the maximum of the lowest 10% of the intensity values and $a_3$ is $\pi/2$, as we expect fiber alignment in the horizontal direction (see Fig. 1a, where $\theta = 0$ corresponds to up-down alignment and $\theta = \pi/2$ to left-right alignment). To obtain the nematic order parameter $S$ from the ODF we start with the expression used in Ref. [1], following a similar procedure as in the derivation starting at

Equation 2, and where $\frac{3}{2}\cos^2(\theta) - \frac{1}{2}$ is used as the form factor of a single rod. We find:

$$S = 2\pi \int_0^\pi \left(\frac{3}{2}\cos^2(\theta) - \frac{1}{2}\right) \Phi(\theta) d\cos(\theta) \tag{13}$$

$$= 2\pi \int_0^\pi \left(\frac{3}{2}\cos^2(\theta) - \frac{1}{2}\right) \frac{\sqrt{m}e^{m\cos(\theta)^2}}{2\pi^{3/2}\mathrm{erfi}(\sqrt{m})} d\cos(\theta) \tag{14}$$

$$= \frac{\sqrt{m}}{2\sqrt{\pi}\,\mathrm{erfi}(\sqrt{m})} \int_0^\pi \left(3\cos^2(\theta) - 1\right) e^{m\cos(\theta)^2} d\cos(\theta) \tag{15}$$

$$= \frac{1}{2\sqrt{\pi}\,\mathrm{erfi}(\sqrt{m})} \int_0^\pi \left(\frac{3}{m}(\sqrt{m}\cos(\theta))^2 - 1\right) e^{(\sqrt{m}\cos(\theta))^2} d\sqrt{m}\cos(\theta) \tag{16}$$

$$= \frac{1}{2\sqrt{\pi}\,\mathrm{erfi}(\sqrt{m})} \int_{-\sqrt{m}}^{\sqrt{m}} \left(\frac{3}{m}x^2 - 1\right) e^{x^2} dx \tag{17}$$

$$= \frac{1}{2\sqrt{\pi}\,\mathrm{erfi}(\sqrt{m})} \left(\int_{-\sqrt{m}}^{\sqrt{m}} \frac{3}{m}x^2e^{x^2}dx - \int_{-\sqrt{m}}^{\sqrt{m}} e^{x^2}dx\right) \tag{18}$$

$$= \frac{1}{2\sqrt{\pi}\,\mathrm{erfi}(\sqrt{m})} \left(\int_{-\sqrt{m}}^{\sqrt{m}} \frac{3}{m}x^2e^{x^2}dx - \sqrt{\pi}\,\mathrm{erfi}(\sqrt{m})\right) \tag{19}$$

To integrate $\int_{-\sqrt{m}}^{\sqrt{m}} x^2e^{x^2}dx$ we use integration by parts ($\int u\,dv = u\,v - \int v\,du$):

$$\int_{-\sqrt{m}}^{\sqrt{m}} x^2e^{x^2}dx \tag{20}$$

$$= \frac{1}{2}\int_{-\sqrt{m}}^{\sqrt{m}} x(2xe^{x^2}dx) \tag{21}$$

$$= \frac{1}{2}x\,e^{x^2}\|_{-\sqrt{m}}^{\sqrt{m}} - \frac{1}{2}\int_{-\sqrt{m}}^{\sqrt{m}} e^{x^2}dx \tag{22}$$

$$= \sqrt{m}\,e^m - \frac{1}{2}\sqrt{\pi}\,\mathrm{erfi}(\sqrt{m}) \tag{23}$$

so that the expression for $S$ becomes

$$S = \frac{1}{2\sqrt{\pi}\,\mathrm{erfi}(\sqrt{m})} \left(\int_{-\sqrt{m}}^{\sqrt{m}} \frac{3}{m}x^2e^{x^2}dx - \sqrt{\pi}\,\mathrm{erfi}(\sqrt{m})\right) \tag{24}$$

$$= \frac{1}{2\sqrt{\pi}\,\mathrm{erfi}(\sqrt{m})} \left(\frac{3}{m}\sqrt{m}\,e^m - \frac{3}{2m}\sqrt{\pi}\,\mathrm{erfi}(\sqrt{m}) - \sqrt{\pi}\,\mathrm{erfi}(\sqrt{m})\right) \tag{25}$$

$$= 2\pi\left(\frac{3e^m}{4\sqrt{m}\pi^{3/2}\mathrm{erfi}(\sqrt{m})} - \frac{3+2m}{8m\pi}\right) \tag{26}$$
